# Seedless: On-the-fly pulse calculation for NMR experiments

**DOI:** 10.1101/2024.01.31.578133

**Authors:** Charles Buchanan, Gaurav Bhole, Gogulan Karunanithy, Virginia Casablancas-Antràs, Adeline Poh, Benjamin G. Davis, Jonathan A. Jones, Andrew J. Baldwin

## Abstract

All NMR experiments require sequences of RF ‘pulses’ to manipulate nuclear spins. Signal is lost due to non-uniform excitation of nuclear spins resonating at different energies (chemical shifts) and inhomogeneity in the RF actually generated by hardware over the sample volume. Here, we present Seedless, a tool to calculate NMR pulses that compensate for these effects to enhance control of magnetisation and boost signal. The calculations take only a few seconds using an optimised GRadient Ascent Pulse Engineering (GRAPE) implementation, allowing pulses to be generated ‘on-the-fly’, optimised for individual samples and spectrometers. Each calculated pulse requires bands of chemical shift to be identified, over which one of 4 transforms will be performed, selected from a set that covers all commonly used applications: a universal rotation (e.g. 90° about the x axis), including as a special case an identity operation (return spins in the same state as they started), state-to-state (e.g. Z->Y), an XYcite (Z->XY plane), or a novel type, a suppression, that leaves spins minimally perturbed at all times during the pulse. Using imaging experiments we demonstrate our pulses effectively increase the size of the coil volume and effectively increase signal-to-noise in all experiments. We illustrate the approach by devising ultra-broadband pulses (300 ppm excitation pulse for ^19^F 1D spectra), a ^15^N HSQC with 58% increased S/N (950 MHz spectrometer + cryoprobe), triple resonance biomolecular NMR experiments such as HNCACO with 55% increased S/N (600 MHz spectrometer + RT probe), and a highly efficient pulse sequence for water suppression. The 8 optimised pulse sequences presented required 54 bespoke pulses all calculated on-the-fly. Seedless provides a means to enhance sensitivity in all pulse sequences in a manner that can be tailored to all samples/hardware being used.

## Introduction

Nuclear magnetic resonance spectroscopy is one of the most widely used techniques for atomic resolution characterisation of molecules and biomolecules. NMR experiments, termed pulse sequences, can be selected from vast libraries each providing different molecular characterisations. These are (predominantly) sequences of radio-frequency (RF) ‘pulses’ and delays that together conduct specific manipulations of the nuclear spins in the sample. Pulses are themselves constructed from a series of concatenated ‘rectangular’ elements, each of which has a defined ‘phase’ (angle in the xy plane), a central frequency (transition energy), amplitude (intensity), and duration. The simplest ‘rectangular’ pulses have constant phase, frequency and amplitude.

Maximum sensitivity in experiments requires pulses that have a uniform performance both spatially (all positions in the sample experience the same excitation), and spectroscopically (where the effects of the pulse are identical over some range of chemical shifts). In practical situations neither of these conditions are met. Spatially, a rectangular pulse will have a variation in amplitude of approximately ±5% variation over the sample from the desired value, with the specific values varying with the hardware^1^. Spectroscopically, a rectangular pulse will have an excitation profile described by a sinc function where ‘perfect’ excitation occurs only for nuclear spins whose transition energy is close to the central frequency. As spectrometers are constructed with higher fields, when examining samples that contain a wide range of chemical shifts, and when performing experiments with larger numbers of pulses, these problems become more acute and sensitivity is lost.

To address this, many ‘shaped’ pulses have been developed^2-7^, comprising concatenated rectangular elements each of which can have varying amplitude and phase, which can be collectively optimised to produce specific actions such as inversion, excitation or refocusing^2-7^ or for wide broadband excitation^8^. The present paradigm remains the same as first introduced for the BURP pulses in 1991^6^, which is to calculate an optimised ‘shape’, with desirable characteristics, where total duration/amplitude can be rescaled to fit specific applications at the point of application. The field received a significant boost with development of GRadient Ascent Pulse Engineering (GRAPE) methods, which allow efficient calculation of the derivatives required for efficient optimisation^9^. Implementations of the GRAPE algorithm typically require proprietary software such as Spinach^10^ that requires MATLAB, or are embedded in general frameworks not specifically optimised for performance such as QuTIP^11^ and SIMPSON^12^. Using these tools an impressive range of bespoke pulses have been calculated and made available for use by the community^13-20^. More recently AI algorithms have also been trained for this purpose^21,22^. However the calculation of each pulse can still take many hours, and sequences are still constructed from a small library of underlying designs. Moreover as different buffers alter the fields required for the desired transformations, the excitation profile will change possibly compromising performance.

To move beyond this paradigm, it is highly desirable to calculate bespoke pulses on-the-fly within a few seconds at an NMR spectrometer, where multiple sets of requirements can be matched to a specific sample/experiment/spectrometer. To address this we have developed Seedless (GRAPE without the seeds), highly efficient open-source software written in C++ that can be easily compiled in windows, linux and mac operating systems. Using Seedless, the requirements for each pulse including specifications for chemical shift ranges to perform specific rotations can in principle be directly stored in a pulse sequence, and all required pulses can be calculated in a bespoke fashion within seconds at the point an experiment is started. We either match or exceed the expected performance of existing GRAPE pulses (**section S6**).

The performance of the Seedless algorithm relies on an efficient implementation of the GRAPE algorithm for isolated spin ½ nuclei that dramatically reduces the number of calculations that need to be performed when optimising pulses (derivations in **section S2**). In practical applications, a nucleus, peak field strength, amplitude (peak B_1_ field), duration, carrier frequency (in ppm) and number of segments are specified, together with one or more ppm ranges (bands) each aiming to performing specific transforms (**table 1**, usage instructions **section S5**). In general, the more ‘demanding’ the restraints on the pulse as defined by the pulse sequence and the number of distinct ‘bands’ in chemical shift space, the longer the pulse needs to be, and we discuss principles to follow when selecting transforms to optimise different applications (**section S3**). The pulses generated are compensated for amplitude inhomogeneity by a simple and efficient method, which can be tailored to a measured distribution required for a specific probe. We demonstrate via NMR imaging experiments that the effect of this is to increase the coil volume of the spectrometer, dramatically increasing the signal to noise (**Fig. 1A**).

**Table 1:** A summary of the 4 restraint types that can be applied to nuclear spins in a Seedless calculation, together with the effective number of axes are being controlled (a detailed derived is provided **section S2**). To control more axes, either a longer total duration is required or a higher maximum amplitude. A commonframework is exploited in seedless, where the overall propagator *V* and a related quantity for gradients *C*_*j*_ (see text) are first calculated looping over all elements of the pulse, followed by evaluation of W, A, and B, together with the infidelity ‘cost’ function I. A final loop over all elements is then required to calculate the derivatives (final column) which are used for optimisation. A universal rotation requires either an axis and angle to be supplied (eg 90_o_ x), or the identity operator. A state-to-state pulse requires starting and finishing matrices state *ρ*_*s*_ and *ρf* to be supplied (as X, Y or Z). The XYcite restraint follows a different form, as here we are aiming to select ‘against’ having a Z component by the end of the pulse, rather than, as in all other cases, selecting ‘for’ a state. The ‘suppression’ restraint effectively repeats a state-to-state infidelity evaluation for each time point during the pulse and so is a relatively demanding computation.

| Type | Axes | $W$ | $A$ | $I$ | $B$ | $\frac{dI}{d\phi_j}$ |
| --- | --- | --- | --- | --- | --- | --- |
| **Universal/Identity**<br>(**section S2.4**) | 3 | $\frac{1}{2}U^\dagger$ | $VW$ | $1 - \text{Tr}(A)$ | $-A$ | $\text{Re}[\text{Tr}(BC_j)]$ |
| **State-to-state**<br>(**section S2.2**) | 1 | $\rho_s V^\dagger \rho_f$ | $VW$ | $1 - \text{Tr}(A)$ | $-2A$ | $\text{Re}[\text{Tr}(BC_j)]$ |
| **XYcite**<br>(**section S2.6**) | <1 | $\rho_z V^\dagger \rho_z$ | $VW$ | $[\text{Tr}(A)]^2$ | $4\text{Tr}(A)A$ | $\text{Re}[\text{Tr}(BC_j)]$ |
| **Suppression**<br>(**section S2.7**) | 1 | $\rho_z X_k^\dagger \rho_z$ | $X_k W_K$ | $1 - \sum_{k=1}^n \text{Tr}(A_k)$ | $-2A_k$ | $\sum_{k=j}^n \text{Re}[\text{Tr}(B_k C_j)]$ |

**Figure 1:**
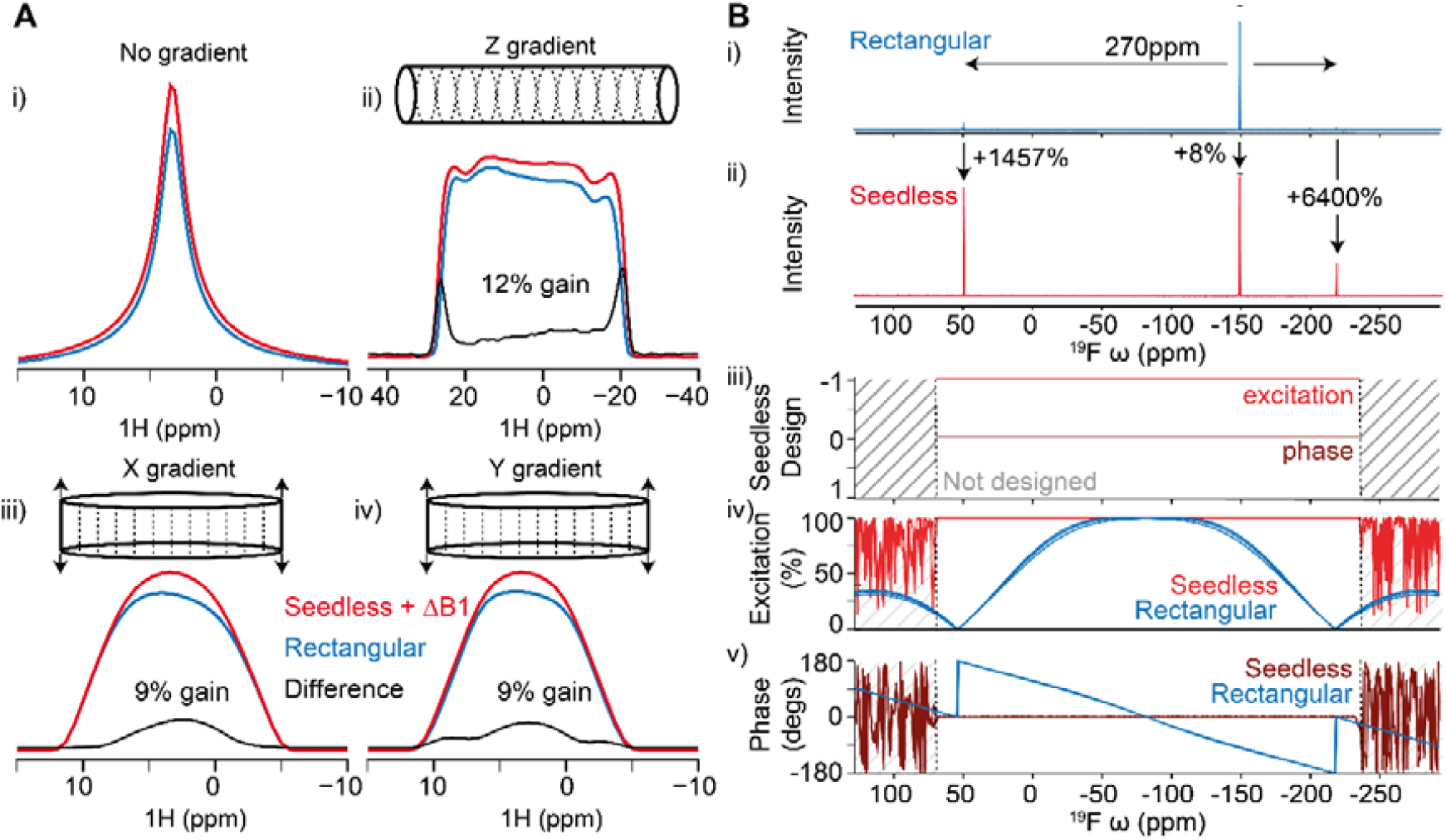
**(A)** A^13^ C HSQC imaging gradient echo sequence was designed (**section S3.2.1**) and implemented with both seedless (red) and rectangular (blue) pulses on a sample of ^13^C methanol on a probe equipped with XYZ gradients. (**Ai**) In the absence of the imaging gradients, the sequence using seedless pulses delivered substantial gains (12% increase in integral), in a case where all pulses were held precisely on-resonance. (**Aii/iii/iv**) the Z, X and Y projected images were obtained allowing the enhanced sensitivity to be attributed to regions of space above the top and below the bottom of the sample, at the centre of the tube. This is precisely where significant divergence is expected in the RF fields generated by the probe. Similar results were obtained for the Z axis when performing the experiment on probes equipped only with Z gradients (**Supplementary Fig. S2**), demonstrating that compensating for B_1_ inhomogeneity can lead to substantial sensitivity improvements by effectively increasing the coil volume. (**Bi**) To test the ability of seedless to construct an ultra-broadband excitation pulse (**section S.3.2.2**), a ^19^F NMR sample was created from Selectfluor and FmocOSu, which together contain 3 F environments that span 270 ppm (**section S.4.2**). Using a 12.6 μs 20 kHz rectangular excitation pulse placed in the average location, allowed only the central resonances to be appreciably discerned (**Supplementary Fig. S3**). A 2 ms seedless excitation pulse (Z-> -Y) at 20.2 kHz was designed and the resulting spectrum (ii) contained all three expected resonances, effectively granting an in increase in intensity of the resonances at the edges of the spectrum by a factor of 1457% and 6400% versus the rectangular pulse. While all resonances obtained in the seedless experiment could be phased using 0 order correction, resonances from the rectangular experiment could not. The simulated profiles for the performance of the rectangular and seedless pulses (**iv**,**v**) are shown for the three different B_1_ amplitude values of 0.95, 1.00 and 1.03 of the peak value. As the individual traces cannot be readily discerned for the Seedless pulse in this figure, the pulse can be considered highly tolerant of amplitude variation.

We demonstrate the effectiveness of seedless by generating a high bandwidth (300 ppm) pulse for quantitative 1D ^19^F spectroscopy (**Fig. 1B**), and using the ‘suppress’ restraint, we created a ‘perfect echo’^23^ 1D pulse sequence that gives a protein 1D NMR spectrum on dilute samples with water artefacts reduced to ca. 7.5 μM levels, a reduction factor > 10^7^ (**Fig 2A**). We then generate a ^15^N HSQC spectrum with peak intensities enhanced by a factor of 58% on spectrometer operating at 950 MHz ^1^H Larmor frequency. We finally calculate a series of pulses suitable for triple resonance biological NMR applications where in ^13^C, we exploit independent control of CO, Cα and Cβ groups to generate triple resonance pulse sequences (HNCACO, HNCO, HNCA, HNCOCA) that boost signal to noise and discuss the principles required to optimise such sequences using seedless. Notably, strategies implemented in pulse sequences to compensate for pulse imperfections can simply be removed when using seedless pulses, simplifying the implementation and yielding NMR spectra that require only 0^th^ order phase corrections with perfect baselines. Overall, the 8 pulse sequences we optimise here require 54 bespoke seedless pulses (**section S3**), all of which are freely downloadable. All are calculated within a few seconds on a 2021 Macbook pro with a 10 core M1 Pro processor and 16 GB RAM (**supplementary Fig. S4**) and detailed usage instructions are provided (**section S5**).

**Figure 2:**
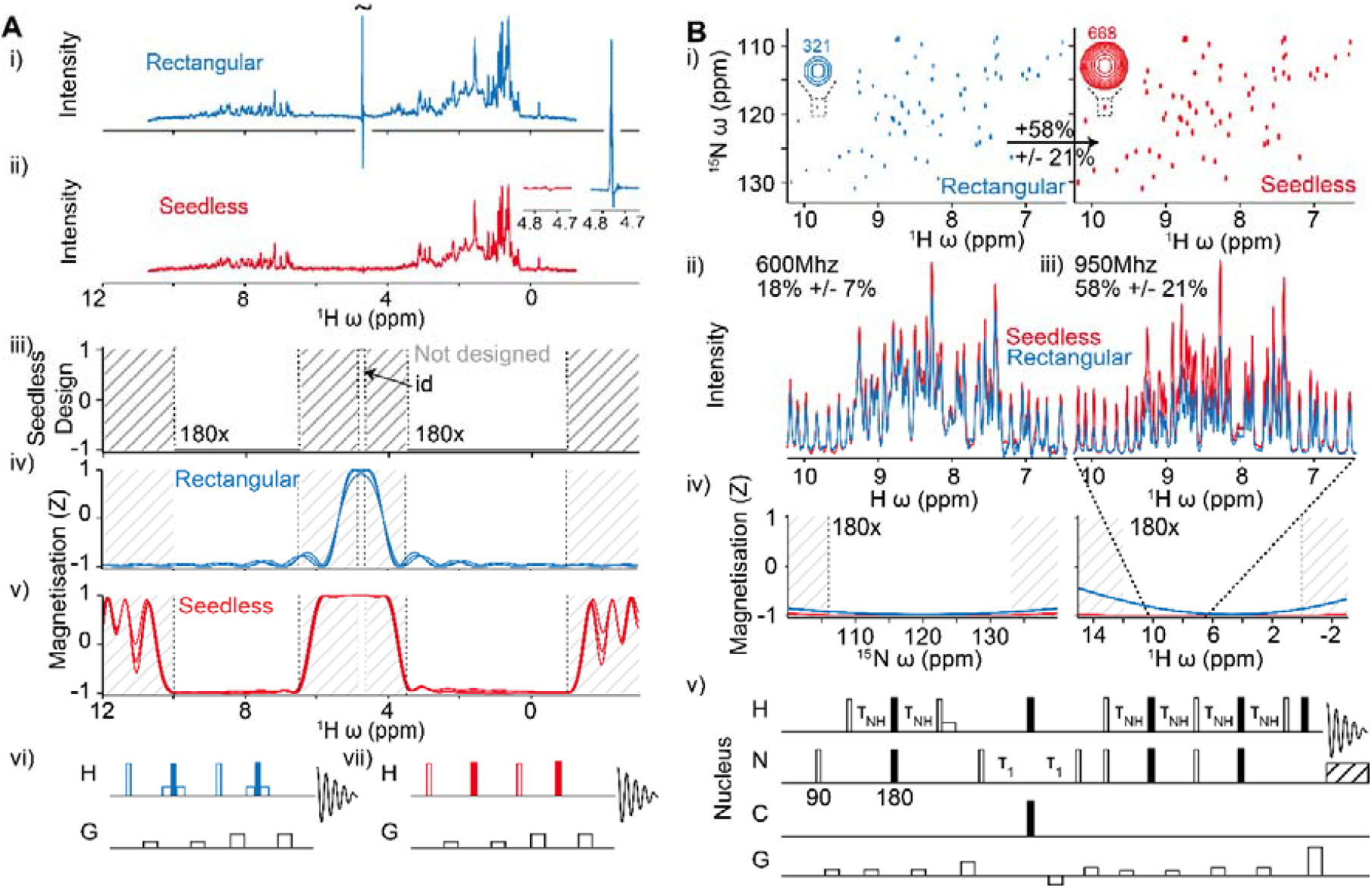
**(A)** A comparison of water-gate perfect echo (PE) pulse sequence using rectangular (**Ai, vi**) and Seedless (**Aii, vii**) pulses (**section S.3.2.3**) (acquired at 600 MHz). For the rectangular pulses, 90 pulses are shown as open boxes while 180 pulses are filled boxes both executed at 24.5 kHz. Lower power (201 Hz), water selective 90 pulses are shown in the rectangular pulse sequence as small boxes. The simulated performance of a rectangular soft/hard/soft 180 composite pulse at these fields is shown (**Aiv**), with the response at amplitudes of 0.95, 1.00 and 1.03 of the central value. The seedless pulses were designed to perform unitary rotations on the aliphatic and amide bands, and suppression of water on the indicated water band (**Aiii**) and the performance of a 4000 μs pulse at 9.76 kHz is shown (**Av, section S3.2.3)**. Spectra were obtained from the two sequences from a sample containing 10 μM lysozyme in PBS with 10% D_2_O (**Ai**,**ii**). Signal originating from water when using rectangular pulses was truncated (tilde). A side-by-side comparison of the residual water signal is shown (inset) with the rectangular sequence given water S/N of 150 and seedless, 2 from a 64 scan (6 minute) experiment (**section S.3.2.3**). The ring shifted methyl protons (S/N 4 in both spectra) allows for an approximate mapping of water signal to an effective concentration, equal to 560 μM/7.5 μM in the rectangular/seedless experiments respectively (suppression factors of 10^5^ and 10^7^). **(B)** An application to ^15^N sensitivity enhanced HSQC spectra (**Bv, section S.3.2.4**), comparing rectangular (blue) and seedless (red) pulses acquired on a cryogenically cooled triple resonance probe operating at 950 MHz (**Bi**) on a sample of U[^1^ H/^13^ C/^15^ N] ABP1P (section S.4.2). Substantial gains are obtained using seedless pulses, which on average were 58% enhanced for each resonance (**Biii**). A strong correlation is obtained of gains versus H chemical shift (**supplementary Fig. S5**). Significant but more modest gains of 18% were obtained on a room temperature probe at 600 MHz (**Bii**), that show no dependence on H chemical shift (**supplementary Fig. S5**).

Seedless allows for pulses to be routinely recalculated with adjusted bandwidths at run-time so that experiments can be optimised for specific samples and spectrometers to boost sensitivity in all pulse sequences, entirely removing the need to store pre-designed ‘shapes’ design for re-use. Seedless is free for academic use and can be downloaded from XXXX.

### Seedless design and implementation

During pulse sequences, each pulse needs to perform specific manipulations of spins that span specified ppm ranges. These can be grouped into one of 4 types of transform (**table 1**), a detailed derivation of each are included in the supplementary information (**section S2**). Here, we describe the key results, and provide principles that can guide decisions for how individual pulses should be replaced with seedless pulses (detailed in **section S3**), before demonstrating specific applications.

All pulses within a sequence aim to take a spin from a starting state (or states) s to a target *f*. If the pulse isn’t perfect, its action will be to create the *f*’. The Uhlmann-Jozsa fidelity 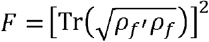 ^24,25^ allows the similarity of two density matrices (*ρ*) describing *f* and *f*’ to be measured^24,25^, varying between 0 (dissimilar) to 1 (identical), the latter being achieved only when the pulse exactly creates the target state from the starting state. For systems of spin ½ all states can be treated as quantum mechanically ‘pure’, and the fidelity simplifies to 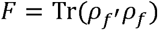. ^24,26^ (**section S.2.4**). The action of a pulse can be mathematically expressed as a time ordered product of *n* element propagators, *V* = *V*_*n*_*V*_*n*-1_ … *v*_2_ *v*_1_, where each *V*_k_ is a unitary transformation matrix describing the action of element *k* (**section S2.2**) each parameterised by an amplitude and a phase, and so 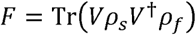. Defining 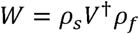, then the fidelity can be written as a product of the pulse, *V*, and a restraint function, W, such that *F* = Tr(*VW*). We use this to construct *I*, the ‘infidelity’ (**Table 1**), a cost function that whose value is equal to 0 when the required action has been achieved.

Such a ‘state-to-state’ (S2S) action will perform transformations such as Z→ −Y, but will not perform the two other cardinal transformations expected from a complete rotation of the Bloch sphere, namely Y→Z and X→X. We can accomplish such a ‘universal’ rotation encompassing all expected transformations by defining *U* as the ‘desired’ unitary transformation propagator and integrating over all possible starting orientations. In this case, the calculation is identical but with 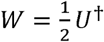 (**Table 1, section S2.4**). A universal rotation restraint could be obtained by applying two S2S restraints on a spin, but our framework accomplishes this at half the cost.

NMR experiments frequently require different transformations to be applied to spins that resonate within distinguishable ‘bands’ of chemical shift. Seedless allows a user to specify as many chemical shift ‘bands’ as required, each with 1 of the 4 types of restraint imposed (**Table 1**). As a general principle, increasing the number of axes of rotation that a pulse needs to control leads to either an increased total duration or maximum amplitude required to retain the same infidelity. Pulses shown in this work are typically operating at the highest possible amplitude allowed unless otherwise stated. A detailed discussion of considerations both general and specific is presented (**section S3**). In brief, the 4 restraints that seedless can impose are:

**1**. A ‘universal’ rotation requires control of all three rotational 3 cardinal axes of the Bloch sphere, independently of any specific starting and finishing state (**section S2.4**). These are the rotations anticipated by an idealised on-resonance rectangular pulse, and performed by the central region of an EBURP1^6^, REBURP^6^ or SURBOP^20^ pulses. Pulses of this type are essential when the pulse is expected to handle a range of incoming states such as the refocusing by a 180° pulse at the centre of a spin-echo, or where magnetisation aligned with two axes need to be simultaneously handled, such as during the refocus INEPT of a sensitivity-enhanced HSQC (**section S.3.2.4**). An axis and an angle need to be supplied to use this restraint (e.g. 90x or 180y, **Table 1, section S2.4**). Similarly, an ‘identity’ restraint can be imposed using the same formalism that returns spins in the same orientation as they started. This can be visualised as a pulse that rotates a spin about the Z-axis backwards by the exact amount it would have otherwise rotated in the XY plane due to chemical shift evolution during the pulse (**section S2.4**). This is critical during indirect evolution periods where we seek to decouple the effects of certain nuclei in a manner that does not interfere with the evolution of the spin of interest (**sections S3.4-8**) yielding spectra requiring zero phase correction in the indirect dimensions. This also simplifies implementations by removing the need to account for undesirable uncontrolled evolution of chemical shift during pulses during delays within sequences.

**2**. While the universal rotation is the most versatile restraint, it is also expensive in terms of total time/amplitude required. If the desired transformation can be simplified to known start and finishing states, e.g. Z→−Y then a ‘state-to-state’ (S2S) transform can be applied (**table 1, section**

**S2.3/4**). As this requires only one effective axis of control, excitation (Z→−Y), de-excitation (Y→Z) and inversion (Z→−Z) pulses will be shorter than their unitary equivalents, the latter being particularly useful during in INEPT transfers to a passive spin or decoupling. These are desirable in cases where either relaxation or hardware restraints favour shorter pulses such as the triple resonance sequences described here.

**3**. If we want to perform a 90° rotation on nuclear spins that either start or finish on Z, but we do not mind what specific phase they end up with in the XY plane, an ‘XYcite’ restraint can be applied. Such a situation arises during INEPT transfers injecting uncertainty into the specific phase of a spin in the XY plane is not important. The PC9 pulses performs in this fashion^4^. Here, the cost function selects for ‘not Z’ as the final state (**section S2.6**). Because they are even less restrictive than state-to-state pulses, these can be shorter, and are used in pairs where the first is a time / phase reversed version of the second^27^.

**4**. Finally, there are cases where we wish to minimally perturb a spin at all stages during a pulse. For such a ‘suppression’, the pulse can be broken into n sub-pulses V_1_, V_2_V_1_, V_3_V_2_V_1_ and so on, and the ‘hold’ state-to-state restraint Z→Z is applied for each (**section S2.7**). This is a highly demanding restraint as it takes a fidelity calculation from scaling with n, to scaling with *n*^*2*^ /2, but is highly suitable for keeping water on the Z axis, avoiding effects of radiation damping by preventing transverse magnetisation build up in the XY plane at any point in time. We use this to construct a ‘perfect echo’ 1D pulse sequence with a water suppression factor of 10^7^ (Fig 2A).

To render the pulses tolerant to amplitude inhomogeneity, we further ensemble average the infidelity over a user specified amplitude distribution. We find that this can be efficiently accomplished on common hardware by sampling the distribution at just three values, at 0.95, 1.00 and 1.03 of the main field strength, weighted 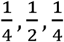 respectively ^1^, reflecting the inhomogeneity distributions measured using a nutation experiment. This can always be measured and adjusted to match any system. We will demonstrate that this alone substantially raises the S/N by effectively increasing the coil volume of a probe (**Fig. 1**).

To generate an optimised pulse, both the phase and the amplitude of each of its n rectangular elements could be optimised to lower the infidelity. The response function of a spectrometer however is not perfect, and when either the phase or amplitude varies suddenly, there will be oscillations around the desired value that eventually dampen out to the target value, typically taking around 100 ns. In this work, to mitigate against this we focused on constant amplitude ‘phase-only’ pulses, with a minimum duration of each step being 2 μs. These are desirable as calculation of the gradients becomes particularly efficient and fewer steps are required for convergence when compared with hybrid amplitude/phase optimisation as has been noted previously^28,29^. Moreover the resulting pulses tend to emerge from calculations in a ‘smooth’ form removing any need to impose additional restraints to enforce this. In all cases explored here, including simultaneous amplitude modulation took a larger number of iterations to arrive at a pulse that wasn’t as good, as judged by the infidelity, as the phase-only variant. Results in this paper are restricted to constant amplitude ‘phase-only’ pulses, analogous to using frequency modulated (FM) radio transmission rather than amplitude modulated (AM).

The Seedless algorithm benefits from two further optimisations (**section S2.8/9**). As we are dealing with calculations for spin ½ nuclei, analytical expressions for the propagators and derivatives have been derived previously^26^ (**section S2.11**). To naïvely evaluate these requires many 2×2 complex matrix multiplications. Because they are ‘scaled-unitary’ (**section S2.12**), we can exploit the symmetry and halve both the number of multiplications required and the memory storage requirements, transforming all matrix/matrix products into vector/vector multiplications. Similarly, taking the trace of the product of two 2×2 matrices should naively require 16 multiplications, but we can account for the ‘scaled-unitary’ symmetry above and reduce this to 4 (**section S2.12**), which particularly accelerates the expensive calculation of the derivatives.

Finally, because all the restraints can be written in the form Tr(*VW*) (**Table 1, section S2**), seedless uses a common framework requiring only two situations where we perform a costly loop over all n pulse elements. In the first loop, the partial products at each step, *xj* =V*j*x*j*-1 are evaluated iteratively, together with a quantity required for the derivatives, 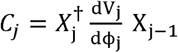 both of which are retained for each element *j* (**section S2.5**). Following this loop, *W, A* and *B* are calculated together with the infidelity cost function *I* (**Table 1**), with the precise form depending on the specific restraint acting on each spin. A final loop over all elements of the pulse calculates the gradients required for optimisation from 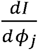 using B and *Cj*. This together enables a highly efficient computation using optimised low level C++ functions. The program makes use of libBFGS, an efficient C++ implementation of the Broyden-Fletcher-Goldfarb-Shanno algorithm^30-33^, which is well suited to this type of optimisation^34^. The code is optionally parallelised using openMP at the level of the two loops over ‘n’, enabling linear rate enhancements with the number of CPU cores provided that the number of spins is sufficiently large that the overhead associated with parallelisation is minimal. The program is controlled via simple input scripts (**section S5**), where lists of pulses can be batch produced. The program optionally produces reports showing the variation in phase/amplitude of the resulting pulses, together with their performance, which was used to generate the figures in this paper and supplementary information.

The resulting package can calculate demanding band selective ^13^C pulses within a few seconds on a 2021 Macbook pro with a 10 core M1 Pro processor and 16 GB RAM. To explore the capabilities of ‘on-the-fly’ pulse calculation, we generated 8 seedless optimised pulse sequences spanning a range of common applications in both chemical, and biomolecular NMR, and conducted detailed experiments to ascertain the precise origins of the enhanced sensitivity.

### Seedless applications

We first considered optimal methods for compensating for the B_1_ field inhomogeneity inherent in modern probes, which can be easily measured using nutation experiments and can be well represented by a distribution containing 3 fields, at 0.95, 1.00 and 1.03 of the main field, weighted at ¼, ½ and ¼ respectively^1^. Starting with a ^13^C enriched methanol sample (**section S4.1**), we performed a 1D ^13^C HSQC spectrum with on-resonance rectangular (**Fig. 1A, blue**), and Seedless pulses (**Fig. 1A red**) with and without amplitude compensation (+/-B_1_), on an NMR spectrometer with a ^1^H Larmor frequency of 600 MHz using a room temperature 5 mm HCN probe equipped with XYZ gradients. The data acquired using the seedless pulses (+B_1_) was 12% more intense than data acquired using rectangular pulses (**Fig. 1Ai**). To visualise specifically which spins were providing additional signal, we adapted a ^13^C/^1^H HSQC imaging pulse sequence to provide Z, X and Y axis projected images (**section S1, S3.2.1**). Taken together, the projections revealed that Seedless pulses were generating more signal predominantly from spins at the top and bottom of the sample, at the centre, which is precisely where we expect the field lines generated by a saddle coil in the probe to show divergence (**Fig 1. Aii**). Similar results were seen when testing on probes equipped only with a Z gradient (**supplementary Fig. S2**). The signal intensity generated from seedless pulses without compensation were comparable to that achieved from a rectangular pulse revealing that the action of the amplitude compensation is to effectively increase the effective coil volume, and so provide enhanced signal intensity. We adopted this compensation strategy in all subsequent pulse designs, which worked equally well on all spectrometer and probe combinations tested in this work (600 and 950 MHz spectrometers with room temperature and cryogenic cooled probes, four combinations in total **section S4.1**).

The total range of ^1^H chemical shifts typically experienced in biological and chemical applications spans ca. 10 ppm, which can be easily excited with a 10 μs rectangular pulse, at an amplitude of 25 kHz, easily generated by modern hardware. By contrast, compounds of interest for ^19^F however can span 300 ppm, which cannot be appreciably excited using rectangular pulses (**Fig. 1B iv**). We sought to design a seedless ultra-wide bandwidth pulse suitable for quantitative analysis^8^. We analysed a sample required for a chemical biology application containing two Selectfluor (containing ^19^F species with chemical shifts 47.89 ppm and -151.5 ppm, **section S4.2**) and a non-natural amino acid analogue FmocOSu (^19^F chemical shift -221.5 ppm). Using a rectangular pulse we could only appreciably discern 1 of the 3 species (**Fig. 1B blue**). We instead designed a 2 ms 25 kHz seedless state-to-state (Z→−Y)excitation pulse designed to span a 300 ppm bandwidth (less than 10 s calculation time, **supplementary Fig. S4**). The Seedless pulse generated an excellent spectrum, with a similar relative intensity ratio of the three species to that obtained from using 3 separate experiments each on resonance with the individual species (**supplementary Fig. S4**). The seedless spectra could be phased with 0^th^ order corrections only, without the need for baseline corrections, giving uniform excitation over the expected range of chemical shifts (**Fig 1Biii, iv, v, Supplementary Fig. S3**). Owing to the excitation profile of the rectangular pulse decreasing sharply at the edge of the spectrum, the signal intensity of elements at the edge of the spectrum was increased by factors exceeding ca. 10^3^ (**Fig 1Bi, ii, Supplementary Fig. S3**).

To further test the applicability of Seedless pulses on more complex biological targets, we prepared a sample of 7 kDa U-[^1^H-^15^N-^13^C] Yeast Actin-Binding Protein 1 (ABP1P, **section S4.2**) as 35 _15_described previousl We first compared a sensitivity enhanced ^15^N HSQC from the Bruker standard library using rectangular pulses to a seedless optimised version where all ^1^H/^13^C/^15^N pulses were replaced with seedless pulses using both a room temperature probe on 600 MHz system and a cryogenically-cooled probe at 950 MHz (detailed description in **section S.3.2.4**). For ease of implementation, all ^1^H/^15^N pulses were unitary rotations, and the decoupling ^13^C 180° pulse was a S2S Z→−inversions requiring 5 seedless pulse designs (**table S.3.1.1**) with total calculation time under 10 s. Pulses were typically 250 μs in duration, at the maximum permitted amplifier power. The corresponding field was determined, on a sample specific fashion from the 90° pulse times, which for ^1^H was 20/23 kHz, for ^15^N was 6.85/7.2 and for ^13^C was 17.6/17.6 kHz at 600/950 MHz respectively.

Individual resonances were interrogated, and at 600 MHz, an average sensitivity gain of 18% was obtained, where the gains were independent of the ^1^H chemical shift (**Fig. 2B, supplementary Fig. S5**) indicating that the benefits arise primarily from amplitude compensation. By contrast, at 950 MHz, the average gains were more substantial, reaching 58% on average (**Fig. 2B**). The gains however were linearly correlated with ^1^H chemical shift (**supplementary Fig. S5**) indicating that both amplitude compensation and improved excitation profiles were together providing the enhanced sensitivity gains.

In principle, the ^1^H and certain ^15^N 90° pulses in the sequence could have been replaced with Shorter Z→− excite/Y→Z de-excite pulses. We note that two 90° pulses in the sensitivity enhanced refocused INEPT period, the first ^15^N and the second ^1^H (**section S.3.2.4**) must perform two simultaneous rotations, either excite/de-excite and the ‘hold’, X→X, operation and so will have to perform universal rotations.

We next sought to design a novel type of pulse motivated by the challenges associated with water suppression. Much of chemistry and biochemistry occurs in water and so it is frequently desirable to generate spectra that allow us to distinguish molecules of interest, as low as ca. 1 μM, from 55 M water. The ‘perfect echo’ (PE) 1D sequence^36^ based on two concatenated ‘watergate’ elements^37^, is an excellent way to do this. Using this sequence, a spectrum containing 10 μM hen egg white lysozyme (HEWL) in PBS with 10% D_2_O yields a water signal with a S/N of 150 in 64 scans (6 minutes). By taking the a ring-shifted methyl proton (S/N=4 from 3 protons at 10 μM) we can estimate an effective water concentration of 560 μM (**Fig. 2A**), and hence assign a suppression factor of 10^5^.

To reduce this contribution, we attempted to directly replace each pulse with Seedless pulses that performed unitary rotations of the aliphatic (-1 to 3.5ppm) and amide (6.5 to 11 ppm) bands, but aimed to leave water not only unperturbed by the end of the pulse, but also to leave water minimally perturbed at the end of all the individual rectangular elements within the pulse (**section S.3.2.3**) and so avoid effects of radiation damping. Using this strategy, the S/N of water was reduced to 2 while leaving the signal in the aliphatic and amide regions unchanged, indicating an effective water concentration of 7.5 μM and a suppression factor of 10^7^ (**Fig. 2A**). The ‘suppression’ pulses were relatively demanding calculations (taking ca. 20 s, **supplementary Fig. S4**). Unlike any other pulse discussed in this manuscript, it was desirable here to use lower amplitudes, ca. 10 kHz to ensure water is minimally perturbed during excitation (**section S3.2.3**). The specific trajectories of water reveal relatively brief periods where water does experience brief excitation (**section S.3.2.3**) but is returned by the pulse to the Z axis to yield excellent protein 1D NMR spectrum at 10 μM with no distortion of the baseline.

Finally, we turned to triple resonance pulse sequences, an important family for biomolecular NMR analysis where correlations and couplings between ^1^H,^15^N, and ^13^C nuclei are exploited^39,40^ to provide atomic resolution characterisation. Sample concentrations of labelled biomolecules are typically lower than those in chemical applications and to accumulate sensitivity, experiment times can extend to days and weeks. We sought to re-design the triple resonance experiments to take advantage of Seedless pulses (**Fig. 3**) to boost sensitivity. This was achieved via several strategies.

**Figure 3:**
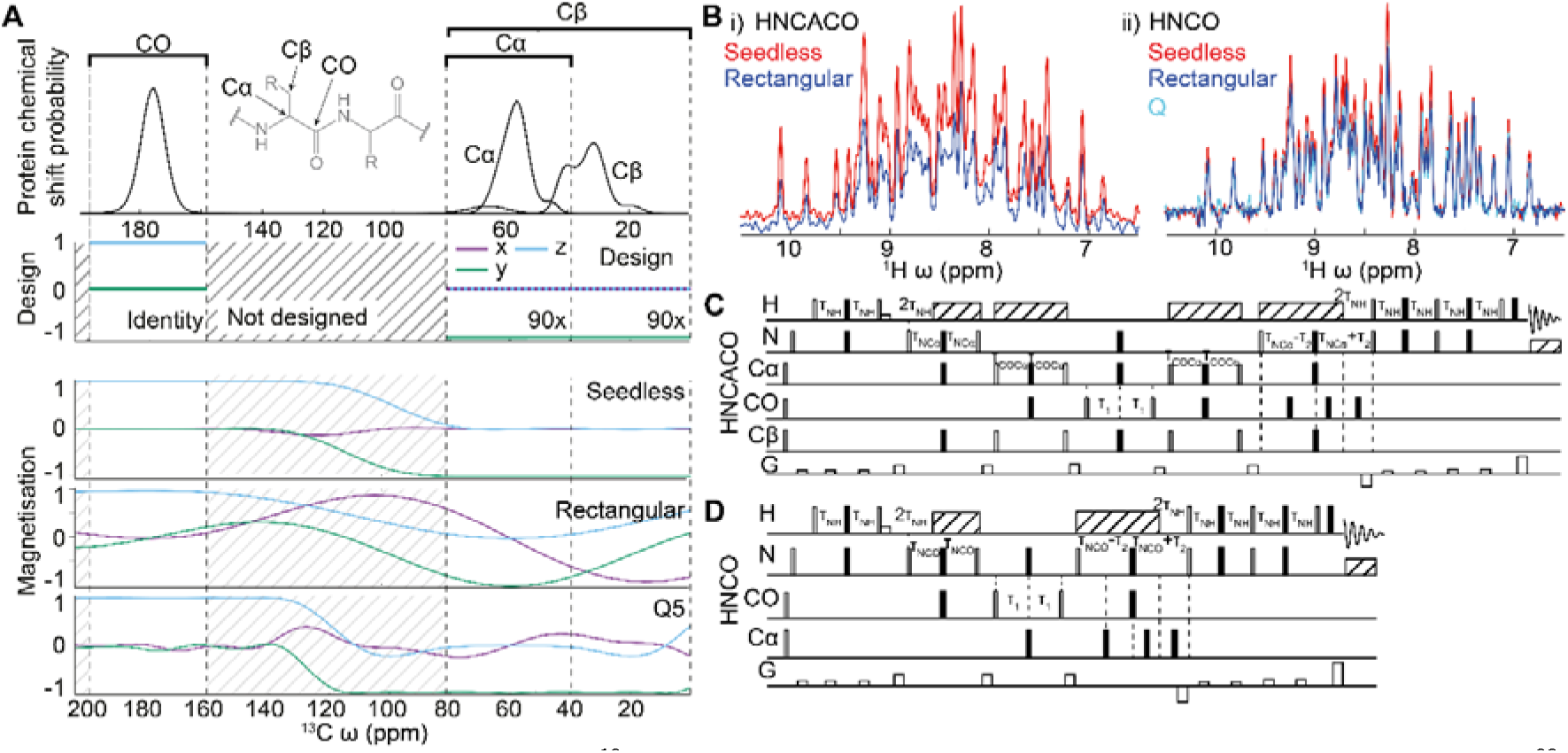
**A i**) The expected distribution of ^13^C chemical shifts expected in proteins, as described by the BMRB. Triple resonance sequences exploit the differences in chemical shift between CO, and Cα/Cβ residues for selective transfers. The overlap of the Cβ and Cα chemical shifts renders it impossible to independently control and hence decouple both groups uniformly for all residues. **A ii**) a simulation of a spin initially aligned on the Z axis following action of a seedless pulse, a rectangular pulse calibrated to perform a 180 rotation at 58 ppm with a first null whose duration is set to 53 μs duration to put the first null at 180 ppm on a 14.4 T spectrometer (**section S3.1**), and a Q3 180 pulse (256 μs at 12.9 kHz with ca. 80 ppm excitation window). Only the seedless pulse please the CO unperturbed and provides a uniform excitation profile of the Cα/Cβ region. (**B**) HN(CA)CO (**C, section S.3.2.7**) and HNCO (**section S.3.2.5**) sequences using rectangular (blue) and seedless (red) pulses were acquired on a U[^1^ H/^13^ C/^15^ N] ABP1P (**D, section S.4.2**). 1D spectra are shown, acquired on a room temperature probe at 600 MHz. Substantial sensitivity gains were obtained using the seedless pulses. The HNCO spectrum was also recorded using Q pulses as implemented in the Bruker standard library, which shows comparable performance to the sequence with rectangular pulses (**Fig. 5**).

The range of chemical shifts spanned by ^13^C in proteins poses both a challenge and an opportunity for triple resonance pulse sequence design as was noted in their original design^41^. Carbonyl (CO) positions resonate in the range 165-185 ppm, and Cα positions fall in the range 40 -65 ppm, where glycine residues (ca. 45 ppm) tend to be the lowest field (**Fig. 3Ai**). The Cβ position is more complicated, with serine and threonine residues resonating in the range 60-90ppm, with the remainder falling in the range 30-50 ppm. Side chain carbon residues then tend to decrease in chemical shift, reaching ca. 10 ppm for the δ methyl carbons of ILE residues (**Fig. 3Ai**). Hardware on high field spectrometers do not allow for uniform excitation from the range 10 ppm to 185 ppm with rectangular pulses (**Fig. 3Aii**), and triple resonance NMR pulse sequences require the different bands (CO, Cα, Cβ) to experience differential excitation, for example, for selective decoupling and magnetisation transfers. Owing to the overlap between the Cα and Cβs, that will depend on the specific sample under study, the two cannot be handled independently in a general case.

Selective excitation was originally accomplished either by using rectangular pulses where the maximum excitation is in the centre of the band of interest and a ‘null’ excitation at the centre of the undesired band (section S3.1), and more recently using selective shaped pulses, such as the Q pulse (e.g. sequences in the Bruker standard library). In both cases, the excitation profiles are not uniform, and spins can evolve during the pulses causing artefacts and loss of signal intensity (Fig 3Aiv). To partially compensate for these effects, additional delays and pulses are introduced within sequences complicating their implementation. To generate a family of ^13^C pulses for triple resonance applications using Seedless, three chemical shift bands for CO, Cα and Cβ were defined, where each band is instructed to provide either a universal rotation, a S2S transform or the identity operation (**table 1, supplementary table S.3.1.1/2**).

We focused on the original four triple resonance experiments which are the most widely used for backbone resonance assignment, the HNCO, HNCA, HN(CA)CO and HN(CO)CA^41^. Detailed pulse sequences and discussions are found in **sections S3.2.5-8**. All sequences are a combination of INEPT-style transfers and evolution periods, where the objective of the pulses is uniform excitation of the required bands, and decoupling of unwanted transfer pathways. Following a process of largely trial and error testing different pulse types in different situations, we established a series of general principles to achieve optimal sensitivity. The speed at which seedless could generate pulses allowed us to reliably screen ca. 50 permutations of each sequence, and the rate at which we could analyse peak intensities in resulting 3D spectra using the UnidecNMR peak picking program^42^ rendered this type of screen feasible.

Owing to the relatively high amplitude of ^1^H pulses (23 kHz) and narrow bandwidth, all these pulses could be both relatively short (100 μs) and perform universal rotations. For optimal sensitivity, owing to the lower amplitudes (6.85 kHz and 17.6 kHz respectively), pulses should be provided for the specific task, and not be ‘overspecified’, and avoiding using longer universal rotations where possible. For phase coherent excitation/de-excitations, S2S Z⟶−Y / Y⟶Z transfers of an acceptable quality were shorter than universal equivalents. For 180° pulses, during INEPT periods and ^15^N evolution, the active (transverse) spin always requires a unitary 180° for refocusing as the pulse needs to correctly transform a range of different incoming phases in the xy plane. Passive spins (held longitudinally) however can either have transfer enabled during an INEPT via an S2S inversion Z⟶Z during which if decoupling is required, a ‘hold’ S2S Z⟶Z can be applied to avoid the need to perform the more expensive unitary identity operation. Exceptions to this minimalist principle are found in the sensitivity enhanced refocused INEPT as described for the HSQC as two pathways need to be controlled requiring universal pulses (**section S.3.2.4**). For CO detection, distortion-less spectrum requires a central pulse that performs a S2S Z⟶Z inversion performed on Cα/Cβ. If an ‘identity’ operation is performed on the CO spin during the inversion, the resulting spectra are distortion free and perfectly phased.

Finally, we considered coupling between the Cα and Cβ. Because of the substantial overlap of the Cα and Cβ chemical shifts (**Fig. 3A**), any attempt to perform different transforms on the Cα and the Cβ will be imperfect, leading to an overlap region where neither Cα nor Cβ are well treated. Because coupling of CO to Cβ is weak, can treat Cα and Cβ as identical, unless one or both is transverse which happens in two specific situations in these sequences. In the HN(CA)CO, there is a Cα⟶CO INEPT transfer where Cα is transverse and can couple to the Cβ resulting in signal loss. To mitigate against this, the central ^13^C pulse needs to perform a unitary 180 rotation of the transverse Cα, a S2S inversion of CO (to allow Cα -CO transfer) and a S2S Z->Z ‘hold’ applied to Cβ for decoupling (**section S.3.2.7**). To achieve acceptable performance, the Cα /Cβ interface was placed 40 ppm. The resulting pulse (400 μs at 17.6 kHz) had an interfacial region of +/-4 ppm (**section S.3.2.6/7**). The interface region could be further reduced if the pulse length, spectrometer field or amplitude were increased. This pulse will erode sensitivity of Cα resonances between 45 and 40 ppm but will completely decouple all Cαs from Cβs where the Cβ was 35 ppm or lower (which automatically excludes all serine and threonine residues, **Fig. 3A**).

Similarly, in the HNCA and HN(CO)CA sequences during Cα indirect chemical shift acquisition, for distortion-less spectra there is a need for a pulse that applies an identity operation on Cα, with inversions on both CO and Cβ for decoupling (**sections S.3.2.6/8**). This was also achieved with a 400 μs pulse 17.6 kHz with considerations allowing us to decouple most amino acid types. We recommend the conventional wisdom of recording spectra at a resolution below which the coupling can be resolved, and so the majority Cβ^23^ decoupling manifests as a sensitivity gain (**supplementary Fig. S6**), though experiments targeting specific residue types could easily be constructed^43^.

The sensitivity of the resulting pulse sequences was substantially improved in all cases, with sensitivity gains of 26% (HNCO Fig. 3B, **Fig. 4**), 29% (HNCA, **Fig. 4**), 58% (HNCACO **Fig. 3B**) and 40% (HNCOCA, **Fig. 4**), acquired at 600 MHz on a room temperature probe, as judged from comparing the intensity of individual resonances versus an equivalent sequence acquired using rectangular pulses.

**Figure 4:**
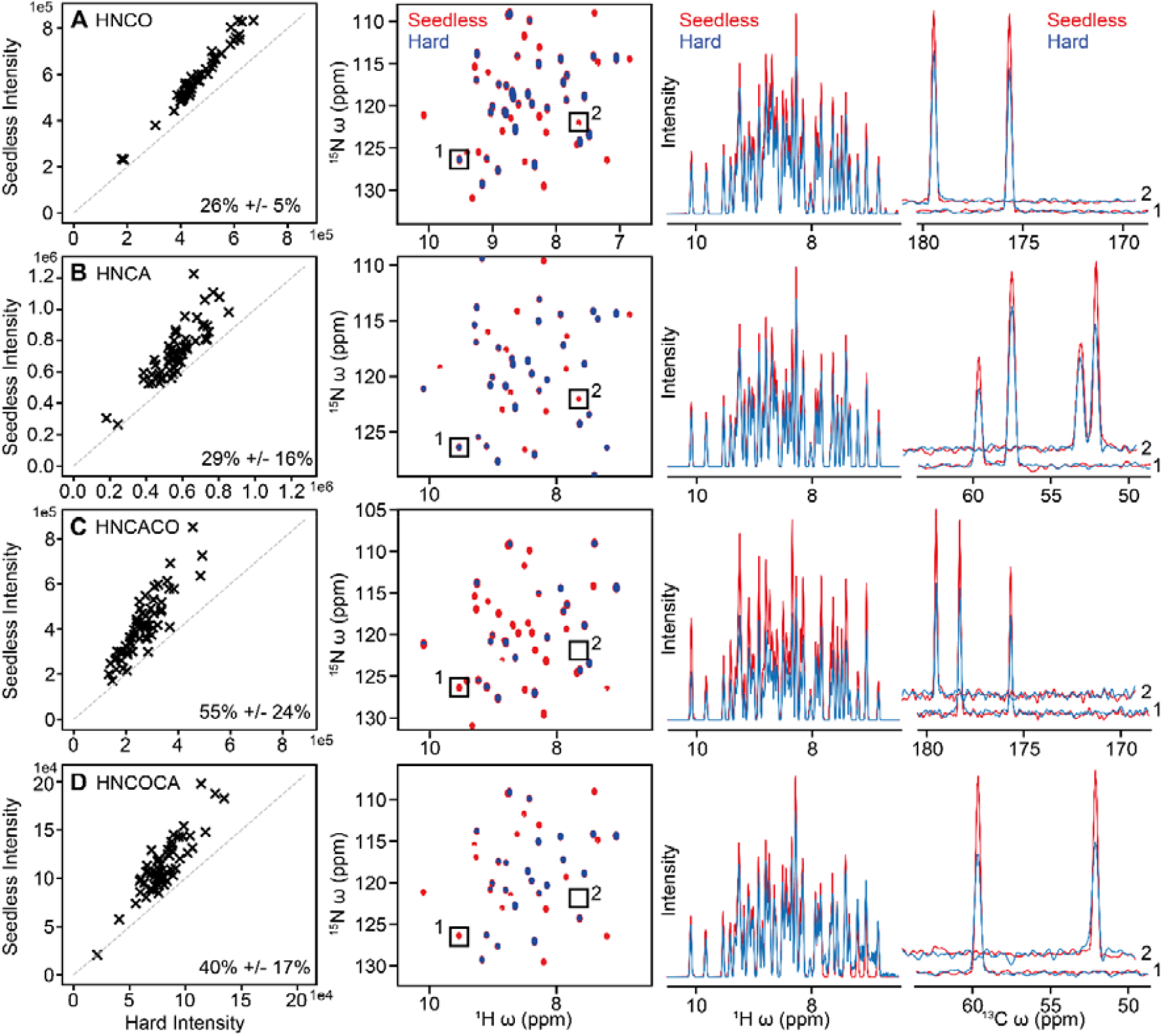
A comparison of signal intensity between seedless optimised and rectangular pulse sequences for HNCO (A), HNCA (B), HNCACO (C) and HNCOCA(D), described in detail in **sections S.3.2.5-8**. ^1^H/^15^ N 2D projections, a 1D H skyline projection and indicates slices (1 and 2) are shown. No peaks were found to have lost intensity upon application of seedless pulses with the intensity of certain residues being doubled.

We sought to understand precisely where our sensitivity gains were coming from and so we carefully deconstructed our treatment of the HNCO. We anticipated correlations with ^13^C chemical shifts, expecting that the improved excitation bandwidth would be largely responsible for our sensitivity gains. This was not the case. The performance of rectangular pulses versus, Q pulses (**Fig. 5ai)** and using only ^13^C seedless pulses without amplitude compensation (**Fig. 5aii**) resulted in similar sensitivities though with noticeable improvements for glycine residues with lower Cα chemical shifts in both cases. By including ^13^C inhomogeneity compensated pulses (**Fig. 5aiii**), then ^15^N (Fig. 5aiv) and then ^1^H (**Fig. 5av**) pulses gave approximately 10% signal to noise gains per modified nucleus channel. On a room temperature probe at 600 MHz therefore, like what was seen for the ^15^N sensitivity enhanced HSQC (**Fig. 3, supplementary Fig. S5**), given no significant correlation between sensitivity improvement and chemical shifts, the gains are coming from effectively increasing the coil volume, where the more pulses in the sequence, the larger the gains. At higher fields, as seen for the ^15^N HSQC, we anticipate further gains coming from both inhomogeneity compensation, and sensitivity enhancements.

**Figure 5:**
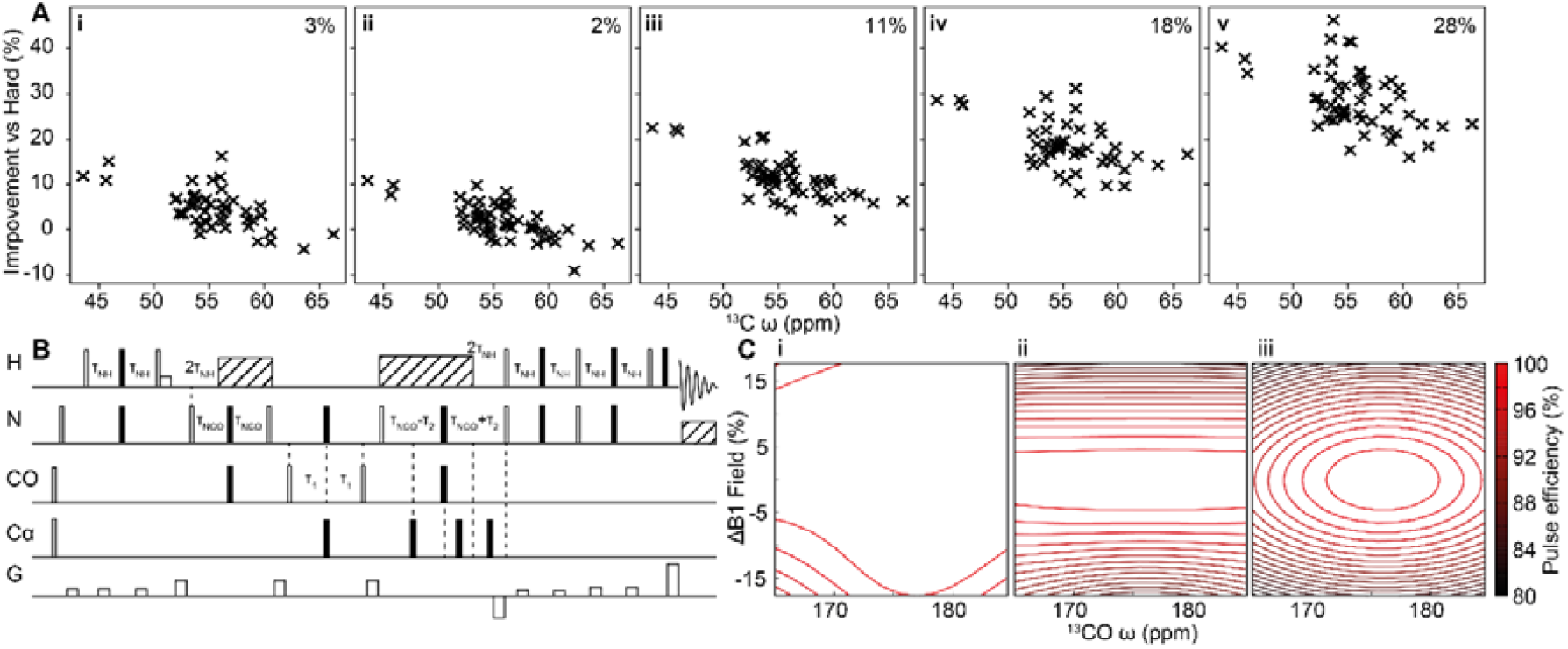
A detailed comparison of the incremental changes made to an HNCO (**section S.3.2.5**) and the variation of sensitivity on Cα chemical shift, as judged on 2D ^13^C/^1^ H planes acquired on U[H/ C/ N] ABP1P on a room temperature probe at 600 MHz. The number stated is the average sensitivity enhancement in each case. (Ai) A pulse sequence from the Bruker standard library using Q pulses rather than rectangular pulses behaved similarly, although glycine peaks (ca. 45 ppm) were appreciably enhanced. (Aii) Non-amplitude compensated ^13^C seedless pulses were included, with sensitivities similar to the sequence with Q pulses. (Aiii) Introducing amplitude compensated C pulses then N (Aiv) and finally ^1^H (Av) yielded approximately 10% increases in S/N with nuclei converted. In this case the sensitivity gains are principally derived from the B_1_ inhomogeneity compensation.

## Discussion

Taken together, Seedless is a versatile tool for calculating pulses for NMR experiment. Owing to an efficient formulation of the theory, pulses with complex restraints can be easily calculated at a rate fast enough for on-the-fly computation. This allows users to easily refine/optimise pulses for specific sample/hardware combinations. Seedless pulses will outperform rectangular pulses and non-homogeneity compensated commonly used shaped pulses such as Q pulses in all sequences and provide a means by which bandwidth can be tuned to the required values. The inhomogeneity compensation effectively increases the coil volume, providing sensitivity increases that scales with the number of pulses in the sequence. In principle, Seedless pulses can be designed with sufficient inhomogeneity correction to be tolerant to the normal fluctuations in sample dielectric, and so removing the need for calibration. Implementing seedless pulses will be far more economical than achieving the same sensitivity gains through purchasing hardware upgrades.

We would in general expect the gains to reflect the specific hardware being used and include factors such as tube geometry. We have tested the pulses at 600 MHz and 950 MHz on both room temperature and cryogenically cooled probes using 5 mm NMR tubes. The inhomogeneity distributions were found to be similar in each case. More substantial gains were seen on 950 MHz spectrometer (58% enhancement of sensitivity enhanced HSQC versus 18% at 600 MHz) reflecting additional substantial gains arising from both improved excitation profiles and homogeneity compensation, with further improvements expected as the field increases further.

We have compared the infidelities of a series of recently produced optimal control pulses against those generated by Seedless under conditions that match (as far as we can ascertain) those originally used. In all cases, seedless pulses achieve lower infidelities though in some cases the difference is modest and would be unlikely to show up as a signal to noise gain at the spectrometer. As seedless pulses are generated in a few seconds and the alternative approaches take many hours, we anticipate that seedless will be a useful general tool for the NMR community.

To create a pulse, a user supplies a peak amplitude (in kHz, determined from a calibrated 90° pulse at the maximum power level), a spectrometer frequency (in MHz), a carrier (in pm), the number of rectangular elements in the pulse, the total duration of the pulse, and a series of bands that span a specified ppm range for a given number of spins, and an operation to be performed. An inhomogeneity distribution is supplied, and the required pulses are returned together (in batch as needed) with a PDF report showing how the pulses perform (**section S5**). Usage instructions are supplied (section S5), and all pulses, pulse sequences and input files associated with this manuscript are available for download. More generally, pulses with a longer duration, and higher amplitude will lead to more effective pulses with sharper response functions between regions of differing chemical shift. In practical applications, the maximum amplitude will be set by hardware restraints, and the maximum duration will be set by the relaxation rates in the system, and so suitable compromises will need to be found. The one exception to this is the water selective ‘suppression’ pulses, where optimisation is required to keep the overall amplitude low and duration (and so to not disturb water during the pulse), while being of sufficient amplitude and duration to have a reasonable boundary between the water band, and the aliphatic and amide excitation bands. Specific and general considerations of how to choose pulses are described both in the text and in the supplementary materials (section S3). The sensitivity of the majority of all routinely used NMR pulse sequences will benefit from the incorporation of these types of pulses.

Seedless is free for academic use. The source code will be available on github, and pre-compiled C++ binaries are available for use under windows, linux and macOS.

## Notes

### Competing Interest Statement

The authors have declared no competing interest.

